# Establishment of a locally adaptive allele in multidimensional continuous space

**DOI:** 10.1101/2024.10.17.618954

**Authors:** Takahiro Sakamoto

**Author notes:** Corresponding author:* National Institute of Genetics, Mishima, Shizuoka, 411-8540, Japan.

## Abstract

Local adaptation is widely seen when species adapt to spatially heterogeneous environments. Although many theoretical studies have investigated the dynamics of local adaptation using two-population models, there remains a need to extend the theoretical framework to continuous space settings, reflecting the real habitats of species. In this study, we use a multidimensional continuous space model and mathematically analyze the establishment process of local adaptation, with a specific emphasis on the relative roles of mutation and migration. First, the role of new mutations is evaluated by deriving the establishment probability of a locally adapted mutation using a branching process and a diffusion approximation. Next, the contribution of immigrants from a neighboring region with similar environmental conditions is considered. Theoretical predictions of the local adaptation rate agreed with the results of Wright–Fisher simulations in both mutation-driven and migration-driven cases. Evolutionary dynamics depend on several factors, including the strength of migration and selection, population density, habitat size, and spatial dimensions. These results offer a theoretical framework for assessing whether mutation or migration predominantly drives convergent local adaptation in spatially continuous environments in the presence of patchy regions with similar environmental conditions.

## Introduction

Many species inhabit spatially varying environments and adapt to local habitats by retaining locally advantageous mutations (Felsenstein 1976; Hereford 2009; Wadgymar *et al*. 2022). The establishment process of these locally adaptive mutations has been a key topic in population genetics theory. Previous studies have often simplified spatial structures into two-population models, successfully describing the evolutionary process of local adaptation. A critical condition for the establishment of locally adaptive alleles is that divergent selection is stronger than migration pressure (Haldane 1930; Felsenstein 1976; Lenormand 2002). Otherwise, one allele dominates the entire population while the other alleles are swamped. Random genetic drift also acts while new mutations are at low frequencies, purging most of them even when they are advantageous. The mechanism by which these evolutionary forces determine establishment probability has been analyzed using a branching process and a diffusion approximation (Barton 1987; Yeaman and Otto 2011; Aeschbacher and Bürger 2014; Yeaman *et al*. 2016; Tomasini and Peischl 2018; Sakamoto and Innan 2019).

Although the establishment process of local adaptation is well understood in the two-population model, its extension to continuous geographical spaces remains incomplete. Considering that almost all species inhabit continuous spaces, it is essential to understand the impact of spatial continuity on the evolutionary process (Bradburd and Ralph 2019). A fundamental distinction between the two-population model and continuous space lies in migration dynamics. The former assumes movement between two locations in a single event, whereas the latter involves migration caused by gradual movements over many generations. This difference changes the evolutionary dynamics in many cases (Battey *et al*. 2020) and also makes the analysis of local adaptation challenging.

Several studies have attempted to employ continuous space models to examine local adaptation; however, these studies have not fully resolved the establishment process. Early studies have focused on equilibrium allele frequencies after successful establishment using deterministic models (Haldane 1948; Fisher 1950; Slatkin 1973; Nagylaki 1975), without considering the stochastic establishment process. To derive establishment probability, Pollak (1966) first developed a branching process framework to consider establishment in an environmentally heterogeneous metapopulation. Barton (1987) extended this framework and derived a general formula for one-dimensional continuous space. Kirkpatrick and Peischl (2013) incorporated temporal fitness fluctuation into the model and discussed the possibility of evolutionary rescue in a one-dimensional space. However, the establishment process in the multidimensional space remains theoretically elusive.

Regarding this issue, the most relevant study was conducted by Ralph and Coop (2015), which examined the relative contributions of new mutations and migration to local adaptation. These authors considered two patchy regions within a multidimensional continuous space where the environment differs from the other regions. Initially, a locally adaptive allele was established in one of patches (the source) but not in the other (the target). Over time, local adaptation evolves in the target patch through either new mutations or immigration from the source patch. Using this model, the authors analyzed the rates of local adaptation driven by mutation and migration. Given that these rates are essentially proportional to establishment probabilities, their research is pertinent to the problem considered in the present study. However, their approximation neglected spatial structure in evaluating the contribution of the new mutations, precluding quantification of how mutations from each geographic position contribute differently to local adaptation. Additionally, their formula for the immigration-driven case includes an undetermined constant and does not provide a precise quantitative prediction. Therefore, although their theory is valuable for estimating evolutionary timescales, a comprehensive quantification of the process is still required.

The present study aims to establish a theoretical framework for understanding local adaptation in *d*-dimensional continuous space (*d* = 1, 2, 3). Although some studies have suggested that the *d* = 3 case lacks biological relevance (Nagylaki 1975), three-dimensional space may exhibit biological importance in certain populations, including somatic cell populations (Fu *et al*. 2022; Noble *et al*. 2022). First, the establishment probability of a locally adaptive mutation is considered. Extending Barton (1987)’s approach to multidimensional scenarios, we derive a differential equation satisfied by the establishment probability and demonstrate how the probability depends on mutation origin and spatial dimensions. Next, based on the model of Ralph and Coop (2015), we investigate acquisition of a locally adaptive allele through migration and derive the waiting time until local adaptation is established. Notably, the current formula contains no undetermined constants, enabling detailed quantitative predictions regarding the rate of local adaptation.

The relative importance of mutation and migration in adaptation to similar environments has been extensively discussed in the context of convergent evolution (Elmer and Meyer 2011; Stern 2013; Rosenblum *et al*. 2014). Both mutation-driven (parallel evolution *in sensu* Stern (2013)) and migration-driven (collateral evolution *in sensu* Stern (2013)) cases are widely observed in empirical systems (Stern 2013; Rosenblum *et al*. 2014). However, the factors determining the relative contributions of each source remain unclear. This study provides a theoretical framework for predicting the dominant mode based on various parameters, including migration strength, selection pressure, habitat size, and distance between habitats.

## Model

A haploid monoecious species distributed at a uniform density *ρ* in *d*-dimensional space (*d* = 1, 2, 3) is considered. Under additive fitness, the following results can be extended to diploid species by substituting *ρ* with 2*ρ*. There exist two distinct environments: environment 1 and environment 2. A biallelic locus A/a is assumed, where allele A is advantageous in environment 1 but deleterious in environment 2. The log fitness of alleles A and a is denoted by *s*_1_ and 0 in environment 1 and −*s*_2_ and 0 in environment 2, respectively (*s*_1_, *s*_2_ > 0). Environmental conditions at any location are assumed to remain constant over time. Throughout this study, a large *ρ* value is assumed to prevent the local fixation of deleterious alleles.

Migration occurs in every generation. In this study, all migration events are short range, and no long-range dispersal is considered. The distance between the offspring position, ***x***, and parent position, ***y***, follows a Gaussian distribution with variance *σ*^2^:

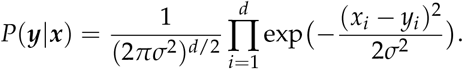

It should be noted that the results may depend on the function of the migration kernel only slightly, provided the moments above the third order of |***x*** − ***y***| are small and the continuous approximation holds, as discussed by Slatkin (1973).

First, we explore local adaptation driven by a new mutation. A target patch with radius *R* is assumed, wherein the interior and exterior environments were designated as environment 1 and environment 2, respectively (Figure 1).

**Figure 1.**
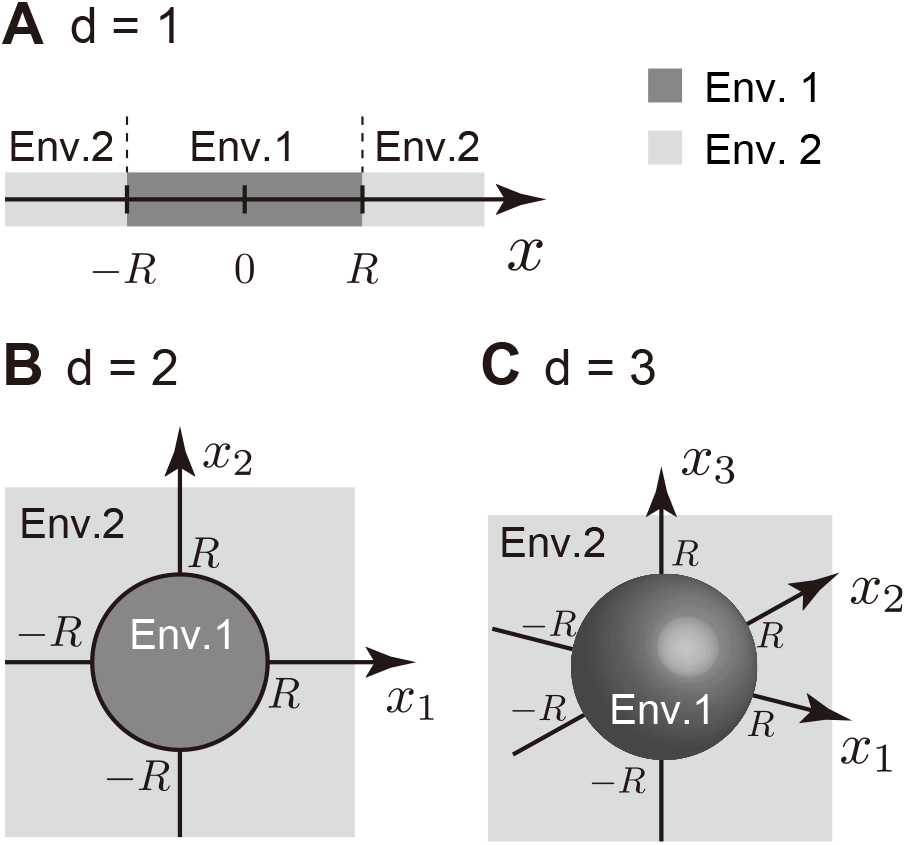
Schematic of the establishment probability model. Patch shapes could be a line segment (*d* = 1) (A), a circle (*d* = 2) (B), and a sphere (*d* = 3) (C).

Depending on *d*, the patch shape could be a line segment (*d* = 1), a circle (*d* = 2), or a sphere (*d* = 3). Initially, allele a is fixed throughout the space and then an allele A arises by mutation. Let *r* denote the distance between the mutation origin and the center of the target patch. If *r* < *R*, the mutation will occur within the target patch, while if *r* > *R*, the mutation will arise in an unsuitable environment. Using this model, the establishment probability of an allele A is investigated. In this study, the establishment probability is defined as the probability that the descendants of an allele reach the target patch, increase in frequency, and are stably maintained at migration-selection balance.

Next, local adaptation via immigration is considered. In this model, another source patch of environment 1 with a radius *R*_*s*_ is assumed (Figure S1). The distance between the source and target patches is denoted by *D*. Initially, allele A is fixed in the source patch, with no allele A outside this patch. The waiting time until allele A is established in the target patch is then considered. In this model, no new mutations arise during the process.

To validate the mathematical analysis, simulations based on the stepping-stone model (Kimura and Weiss 1964) were also conducted. In these simulations, geographic space is discretized into *d*-dimensional grids with migration allowed between adjacent grids. Local adaptation is considered to have occurred when the frequency of allele A within the target patch exceeds 0.5. Details of the simulation settings are provided in APPENDIX A.

## Results

### Establishment probability of a new mutation

The establishment probability of a *de novo* allele is analyzed using a diffusion approximation of a branching process. First, we reviewed the findings of Barton (1987), which studied the establishment probability in one-dimensional continuous space. Subsequently, we extend this framework to the multidimensional case.

#### Overview of Barton (1987)

Barton (1987) considered one-dimensional continuous space, where the position is denoted by *x* (−∞ < *x* < ∞). Initially, allele a is fixed throughout the space. Let *u*_1_(*x*) be the establishment probability of an allele A arising at position *x*. Using the Kolmogorov backward equation, Barton (1987) demonstrated that *u*_1_(*x*) satisfies the following equation:

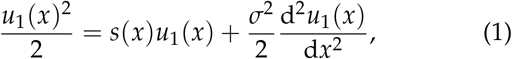

where *s*(*x*) represents the selection coefficient of an allele A at position *x*. The present setting corresponds to the case of *s*(*x*) = *s*_1_ for |*x*| < *R* and *s*(*x*) = −*s*_2_ for |*x*| > *R* (following the local pocket scenario described by Barton (1987)). Barton (1987) derived the solution for *x* > *R* as

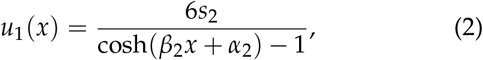

where 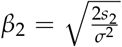 and *α*_2_ is a constant. In this case, *α*_2_ has no explicit formula.

Practically, *α*_2_ can be determined through numerical integration of Equation 1 with boundary conditions 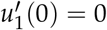 and *u*_1_(∞) = 0. In Figure 2, four sets of *σ*^2^ and *s*_2_ are considered, whereas *s*_1_ = 0.05 and *R* = 1 are fixed. The function *u*_1_ is plotted as black lines against *r* = |*x*|, and vertical dashed lines indicate the boundary between the two environments, *r* = *R*. When a mutation arises in the target patch (*r* < *R*), its establishment probability is ≈ 2*s*_1_, consistent with Haldane’s formula (Haldane 1927). For *r* > *R*, the probability decreases monotonically with increasing *r* (see log-scaled insets in Figure 2). The rate of decrease is primarily determined by *β*_2_ (i.e., the ratio of *s*_2_ and *σ*^2^), where the inverse of *β*_2_ represents the characteristic spatial scale (Slatkin 1973; Ralph and Coop 2015). When *β*_2_*x* is large (i.e., the distance to the target patch is substantially longer than the characteristic spatial scale), establishment of the mutation is almost impossible. In Figure 2, *β*_2_ values are in the panel order A > B = C > D; thus, the rate of decrease is highest in panel A and lowest in panel D, whereas panels B and C exhibit a similar rate, consistent with their *β*_2_ values.

**Figure 2.**
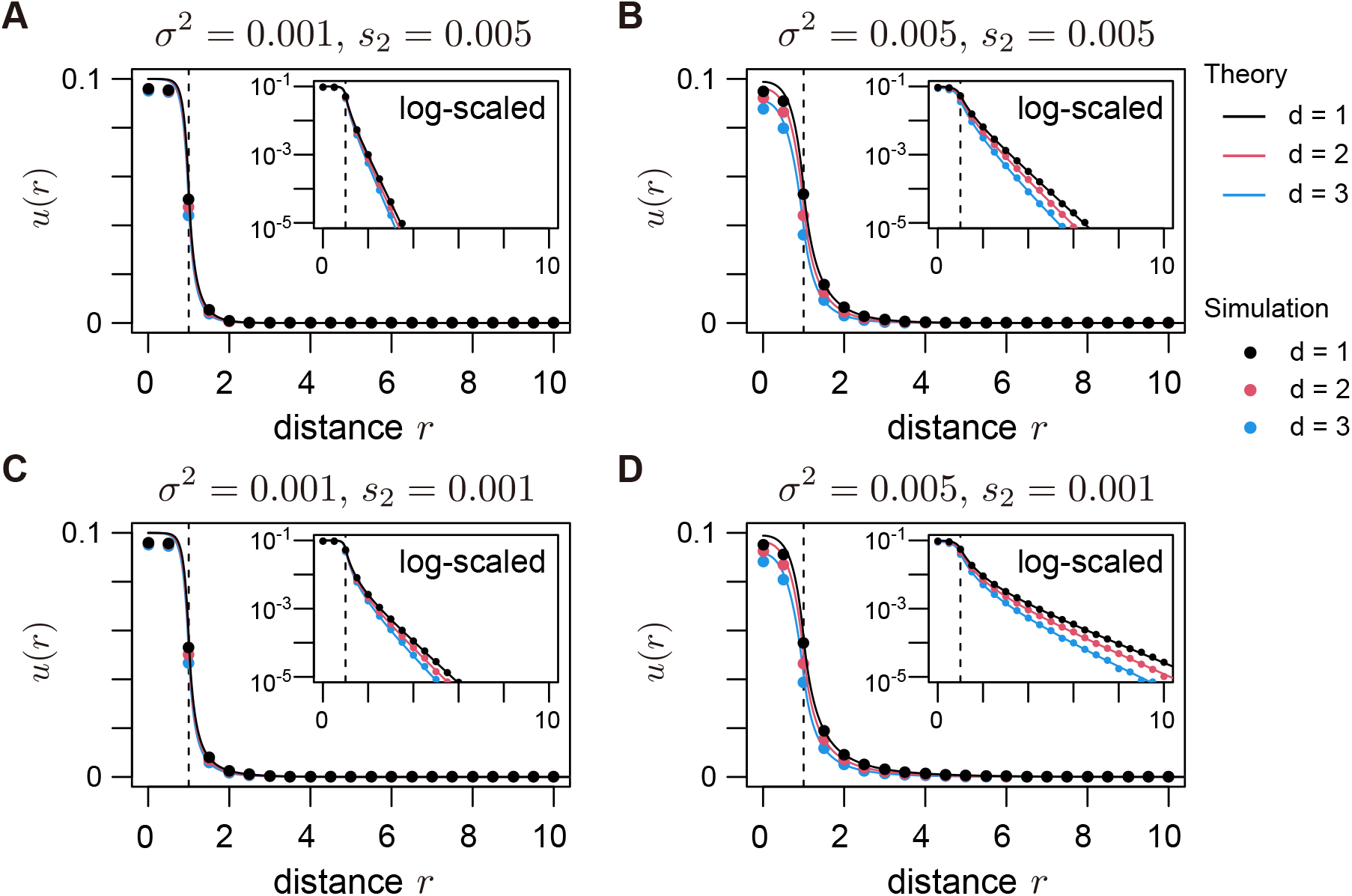
Establishment probability of a locally adaptive mutation as a function of the mutation origin’s position. Four sets of migrational variance and selection were assumed. It was assumed that *s*_1_ = 0.05 and *R* = 1. Solid lines represent theoretical predictions (the solutions to Equations 1 and 3), whereas dots show simulation results. Black, red, and blue represent the results for *d* = 1, 2, and 3, respectively. For each parameter set, 10^7^ simulation replicates were run. Vertical dashed lines are drawn at *r* = *R*, marking the boundary between the two environments.

To validate a diffusion approximation based on a branching process, we conducted simulations with 10^7^ replicates for each parameter set. These simulations assumed a sufficiently large local density of *ρ* = 2^16^ to ensure that local fixation of the deleterious allele is unlikely, as assumed in the branching process approximation. The circles in Figure 2 show agreement between Equation 1 and the simulation results.

#### Extension to the multidimensional case

Equation 1 is extended to derive the establishment probability for *d* ≥ 2. In the *d*-dimensional space, the geographic position is denoted by ***x*** = (*x*_1_, *x*_2_, …, *x*_*d*_). An analog of Equation 1 is given by

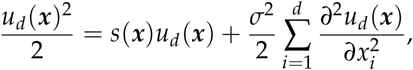

where *s*(***x***) = *s*_1_ for |***x***| < *R* and *s*(***x***) = −*s*_2_ for |***x***| >*R*. Using polar coordinates with *r* = |***x***|, the equation is rewritten:

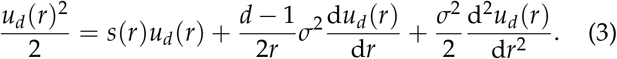

Equation 3 was numerically integrated and *u*_*d*_(*r*) was compared across *d*. Boundary conditions were 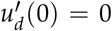 and *u*_*d*_(∞) = 0. In Figure 2, red and blue lines represent *u*_*d*_(*r*) for *d* = 2 and *d* = 3, respectively. The overall trend is similar to that of the *d* = 1 case. When *r* < *R, u*_*d*_(*r*) is approximately 2*s*_1_ and, when *r* > *R, u*_*d*_(*r*) decreases as *r* increases. If *r* is fixed, the establishment probability decreases as *d* increases. This pattern reflects the increased difficulty of reaching the target patch with higher dimensions. Ignoring selection, the allele’s movement is represented by the *d*-dimensional Brownian motion, where the distance to the center of the target patch *r* satisfies the following stochastic differential equation:

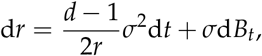

where *B*_*t*_ denotes a standard Brownian motion. Notably, the expected change in *r* in a generation is given by 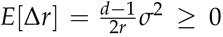, indicating that allele A tends to move away from the center of the target patch for *d* > 1, reducing its establishment probability. This effect strengthens with higher *d* and lower *r*. This effect is also evident in the second term on the right-hand side of Equation 3.

When *r* is large (*β*_2_*r* ≫ 1), *u*_*d*_(*r*) is approximated as follows:

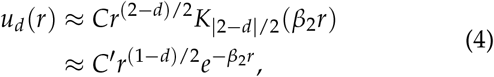

where *K*_*α*_(*x*) is the modified Bessel function of the second kind. This equation is obtained by approximating *u*_*d*_(*r*)^2^ ≈ 0 in Equation 3, as *u*_*d*_(*r*) should be extremely small for large *r*. Equation 4 clearly shows that *u*_*d*_(*r*) decreases more sharply with higher dimensions and larger *β*_2_. The values of *C*s in Equation 4 are determined by the dynamics at small *r* and depend on *σ, s*_1_, *s*_2_, and *d*, although we could not determine them analytically.

We also conducted simulations to check the accuracy of the theory. Equation 3 agrees well with the simulation results (Figure 2), validating the branching process approximation in multidimensional cases as long as the population density (*ρ*) is sufficiently large. In APPENDIX B, we also examine how the simulation results deviate from the theoretical prediction when lower *ρ* is assumed. Because local fixation of allele A reduces the effectiveness of negative selection on allele A, the establishment probability at low population densities could be higher than the theoretical prediction in some parameter sets (Figure B1).

#### Relative probability of the origin distance

What proportion of locally adaptive alleles that are eventually established in the target patch originate at the specific distance from the center of the target patch? In particular, what proportion of such alleles occur within the target patch? To quantify it, the relative contribution of each distance to establishment is calculated as follows:

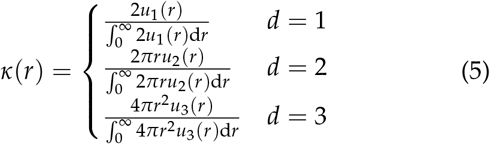

Figure 3 shows the distribution of the origin distances. Each panel corresponds to the parameters used in Figure 2. In the *d* = 1 case (black lines), smaller distances primarily contribute, and most alleles arise within the target patch (*r* < *R*). Conversely, in higher dimensions, the contribution of intermediate distances is more pronounced, as the area at distance *r* increases with *r* in multidimensional cases. When *r* is small, although *u*_*d*_(*r*) is high, the small area results in a relatively lower mutation contribution. When *r* is large, the area becomes larger with many locally adaptive alleles arising; nevertheless, *u*_*d*_(*r*) diminishes markedly, leading to a negligible total contribution. Thus, intermediate distances of *r* ≈ *R* are optimal for producing locally adaptive alleles that establish. Of note, in *d* = 2 and 3 cases, the contribution of the regions outside the target patch may not be negligible especially when *σ*^2^ is large and *s*_2_ is small (Figure 3).

**Figure 3.**
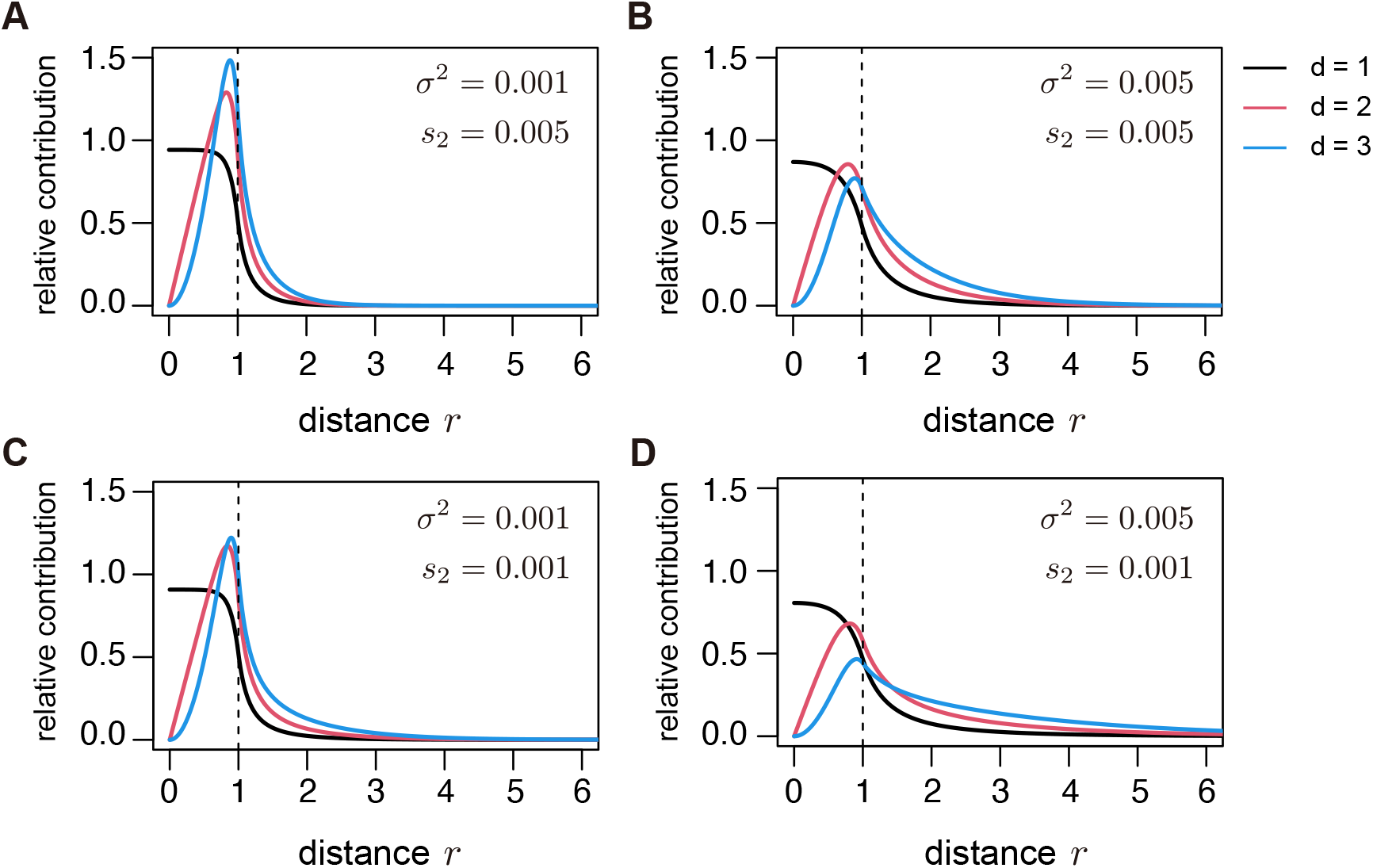
Distribution of the distance between the center of the patch and the arising position of the established allele, *κ*(*r*). Parameter values mirror those given in Figure 2. Vertical dashed lines are drawn at *r* = *R*.

### Immigration from another patch

So far, we have focused on new mutations as a source of locally adaptive alleles. However, another crucial source of local adaptation is immigration from other source patches where locally adaptive alleles have already established (Stern 2013; Ralph and Coop 2015). In this section, we derived an approximation for the waiting time until local adaptation evolves through immigration. Throughout this section, it is assumed that no new mutations arise during the process.

The rate of establishment of a locally adaptive allele driven by immigration is considered the product of the following three factors:

i. The number of allele A leaving the source patch per generation, *N*
ii. The average establishment probability of an allele A that just leaves the source patch, ū_*d*_ (calculated from Equation 3)
iii. The probability that allele A destined to establish does not return to the source patch before reaching the target patch, *P*

Here, *Nū*_*d*_ represents the number of alleles destined to establish under the assumption that the presence of the source patch does not affect the establishment process. The spatial trajectories of these alleles can vary, as shown in Figure 4. Some paths never return to the source patch after leaving (red path in Figure 4), whereas others return by back migration (blue path in Figure 4). We need careful treatment for the latter case. If we simply count the number of emigrating alleles that is destined to be established, the blue allele in Figure 4 is wrongly counted twice because it leaves the source patch twice (before and after back migration). To avoid such overcounting, only alleles that do not return to the source patch should be counted. In this counting scheme, the blue allele is correctly counted once when it was last out of the source patch before the establishment. This procedure necessitates the factor (iii). Since the quantity (ii) was already obtained, the remaining two factors are the focus of this section.

**Figure 4.**
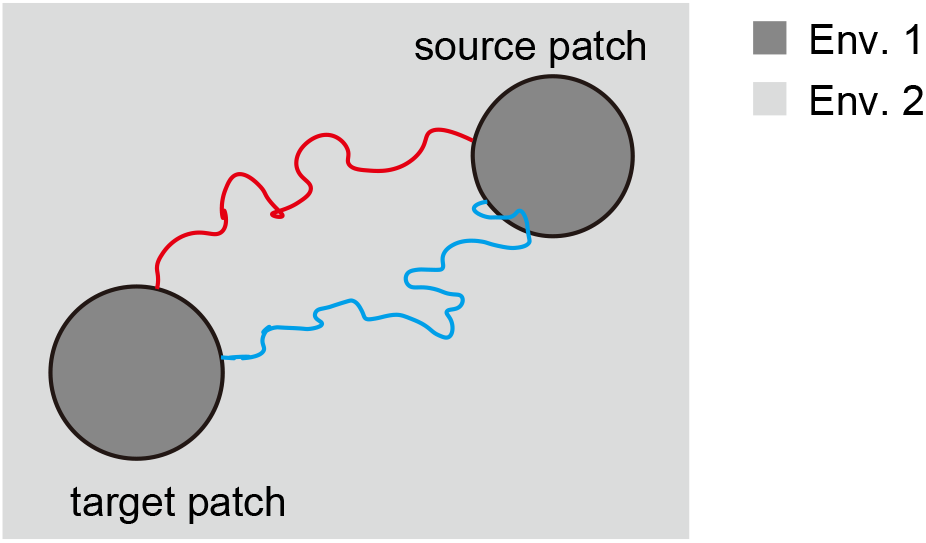
Examples of the geographic trajectories of lineages that eventually establish at the target patch. Red lineage does not return to the source patch after leaving; blue lineage re-enters the source patch before establishment.

#### Outflux rate

First, we consider the number of allele A that emigrate from the source patch in one generation. To derive this, we employed a discrete space approximation. Consider two thin layers surrounding the boundary of the source patch (shown in the enlarged view in Figure 5). One layer is just inside the patch and the other is just outside. The movement from the inner layer to the outer layer is then considered. Let Δ*x* denote the thickness of the layer (Δ*x* ≪ 1/*β*_2_). The number of individuals in the inner layer, *n*(Δ*x*), is given by

**Figure 5.**
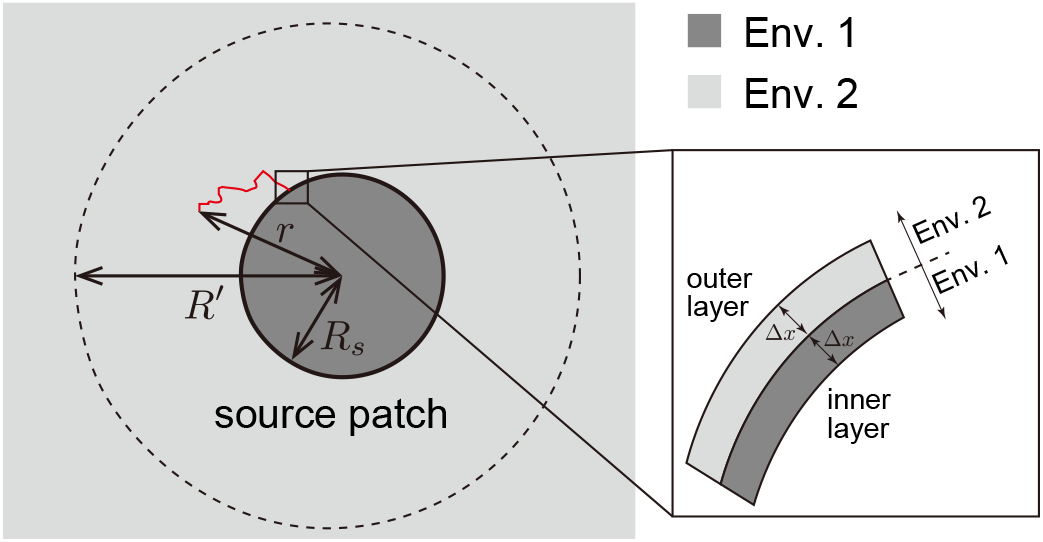
Illustration of the approximations used to calculate *N* and *P*. Enlarged view shows the discrete space approximation at the edge of the source patch. After sufficient time following departure from the source patch, the number of surviving lineages is almost one, depicted by the red path. The stochastic dynamics of this path are analyzed to derive *P*.

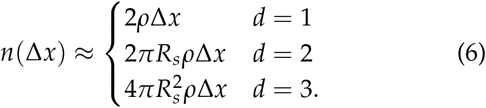

where *R*_*s*_ is the radius of the source patch.

Let *m* be the migration rate between the inner and outer layers in one generation. Note that each migrant from the inner layer moves toward the outer layer or toward the center of the source patch by distance Δ*x*. For the migrational variance to be *σ*^2^, 2*m ×* (Δ*x*)^2^ = *σ*^2^ should be satisfied, resulting in *m* = *σ*^2^/[2(Δ*x*)^2^]. Therefore, the number of allele A leaving the source patch per generation, *N*, is derived as

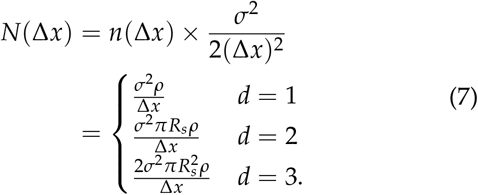

#### Probability of no return

Next, we consider factor (iii). Let us consider the geographic trajectory of offspring alleles of allele A that survive outside the source patch (Figure 5). Due to negative selection, it is unlikely that multiple descendant lineages persist over an extended period and eventually establish at the target patch. Under this scenario, given sufficient time, the number of surviving lineages is almost 0 or 1. The probability that a lineage survives after *t* generations is 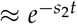; otherwise, all descendants may have gone extinct. In this limiting case, the position of a lineage can by approximated by a *d*-dimensional Brownian motion with a killing process, where the lineage may go extinct at a rate *s*_2_ per generation (see Ralph and Coop 2015, for detailed discussion regarding this approximation).

Using this approximation, we derive the probability that a lineage starting at distance *r* from the center of the source patch reaches a distance *R*^*′*^ (*R*^*′*^ > *r*) before being killed, denoted as *q*_*d*_(*r*) (Figure 5). We begin by allowing the lineage to re-enter the source patch, but such lineages are excluded subsequently. Given that the final focus is on lineages that do not re-enter the source patch, the selection coefficient within the source patch does not affect the final results. Then, for simplicity, we assumed that the log fitness of the lineage was −*s*_2_ throughout the space. Applying the diffusion approximation, *q*_*d*_(*r*) satisfies the following equation:

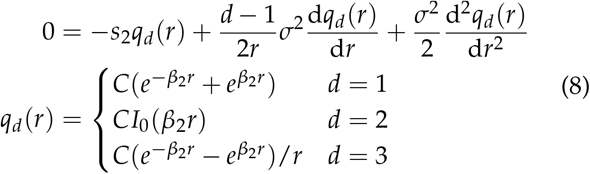

where *I*_*α*_(*x*) is the modified Bessel function of the first kind. The boundary condition 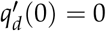 is used, and the constant *C* can be determined through another boundary condition, *q*_*d*_(*R*^*′*^) = 1.

Next, we consider a conditional Brownian motion given that a lineage eventually reaches distance *R*^*′*^. The coefficients of the conditional diffusion process are given by 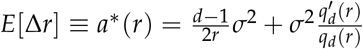 and Var[Δ*r*] ≡ *b*^*^ (*r*) = *σ*^2^ (Ewens 2004). Under this modified diffusion process, the probability that a lineage starting at distance *r* reaches distance *R*^*′*^ before reaching distance *r*_min_ (*r*_min_ < *r* < *R*^*′*^) is calculated as

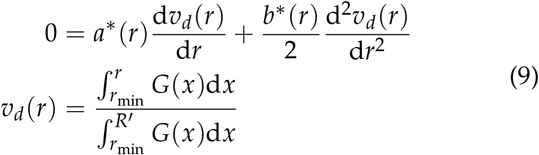

where 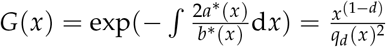.

This result enables the calculation of (iii), assuming *r* = *R*_*s*_ + Δ*x*/2 and *r*_min_ = *R*_*s*_ − Δ*x*/2. The probability that a lineage of allele A at the outer layer does not return to the inner layer before reaching distance *R*^*′*^, conditional on eventually reaching distance *R*^*′*^, is given by

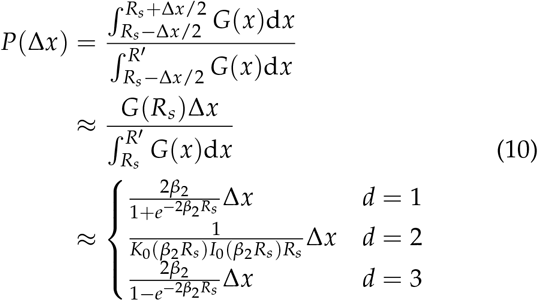

where *K*_*α*_(*x*) is the modified Bessel function of the second kind. In these approximations, we assumed *R*^*′*^≫ 1/*β*_2_ and Δ*x* ≪ 1/*β*_2_. Equation 10 shows that *P*(Δ*x*) is not dependent on *R*^*′*^ if *R*^*′*^ ≫ 1/*β*_2_. This implies that, conditional on reaching the distant target patch, the proportion of lineages that have not re-entered the source patch does not depend on the distance between the target and the source patch.

#### Establishment rate through immigration

Combining Equations 3, 7, and 10, the establishment rate of a locally adaptive allele due to immigration, *λ*_mig_, is given as follows:

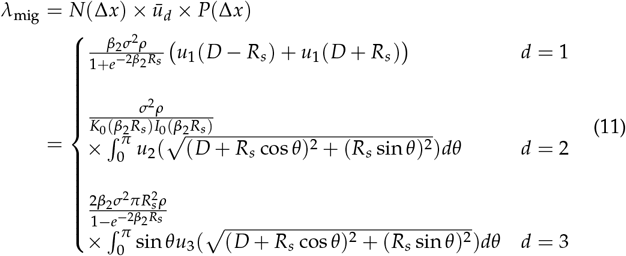

where *D* is the distance between the centers of the target and source patches (Figure S1). In Equation 11, Δ*x* does not appear, indicating that the discretization method does not influence this rate, as expected.

To check the validity of the theory, the expected waiting time 1/*λ*_mig_ is compared with the simulation results. It should be noted that 1/*λ*_mig_ may underestimate the waiting time as it does not consider the time taken by an establishing allele to move from the source patch to the target patch. Ralph and Coop (2015) showed that lineages destined to establish approach the target patch at an average rate of *β*_2_*σ*^2^. Using this approximation, we calculated a modified version incorporating this effect:

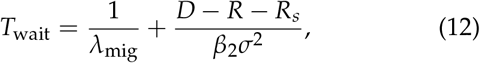

where *D* – *R* − *R*_*s*_ represents the distance between the nearest points of the source and target patches. For comparison, the theoretical prediction of Ralph and Coop (2015) was also calculated based on their Equation 11. Given their equation’s form, *λ*_mig_ = (undetermined constant) *×* (function), we assume the constant to be 1 for simplicity.

The waiting time until mutation establishment through immigrants is shown in Figure 6. In the simulations, 100 replicates were conducted for each parameter set to determine average waiting time. In all cases, *s*_1_ = 0.05, *R* = 1 and *R*_*s*_ = 1 were assumed. Equation 12 agrees well with the simulation results across the four parameter sets investigated. When the distance between the patches is short, a migrant allele destined to establish emerges rapidly; thus, the large proportion of the waiting time is the period required for this allele to reach the target patch. In such a cases, the prediction based on Equation 11 markedly underestimates the waiting time; however, the adjustment made in Equation 12 appears effective. Conversely, when *D* is large, the rate-limiting process is the emergence of a migrant allele that ultimately establishes, and Equation 11 (and Equation 12) agree well with the simulation results. In all cases, the logarithm of waiting time linearly correlated with *D* for large *D*, indicating exponential increases in waiting time as distance increases. Thus, only patches located sufficiently close to the target patch contribute to local adaptation. The slope is mainly determined by *β*_2_ (i.e., the characteristic spatial scale), with the slope’s steepness ranking as A>B ≈ C>D in Figure 6. The theory presented here is consistent with that of Ralph and Coop (2015) (dotted lines), predicting approximately the same slope for large *D*, although the present theory predicts intercepts and offers a more precise quantitative description.

**Figure 6.**
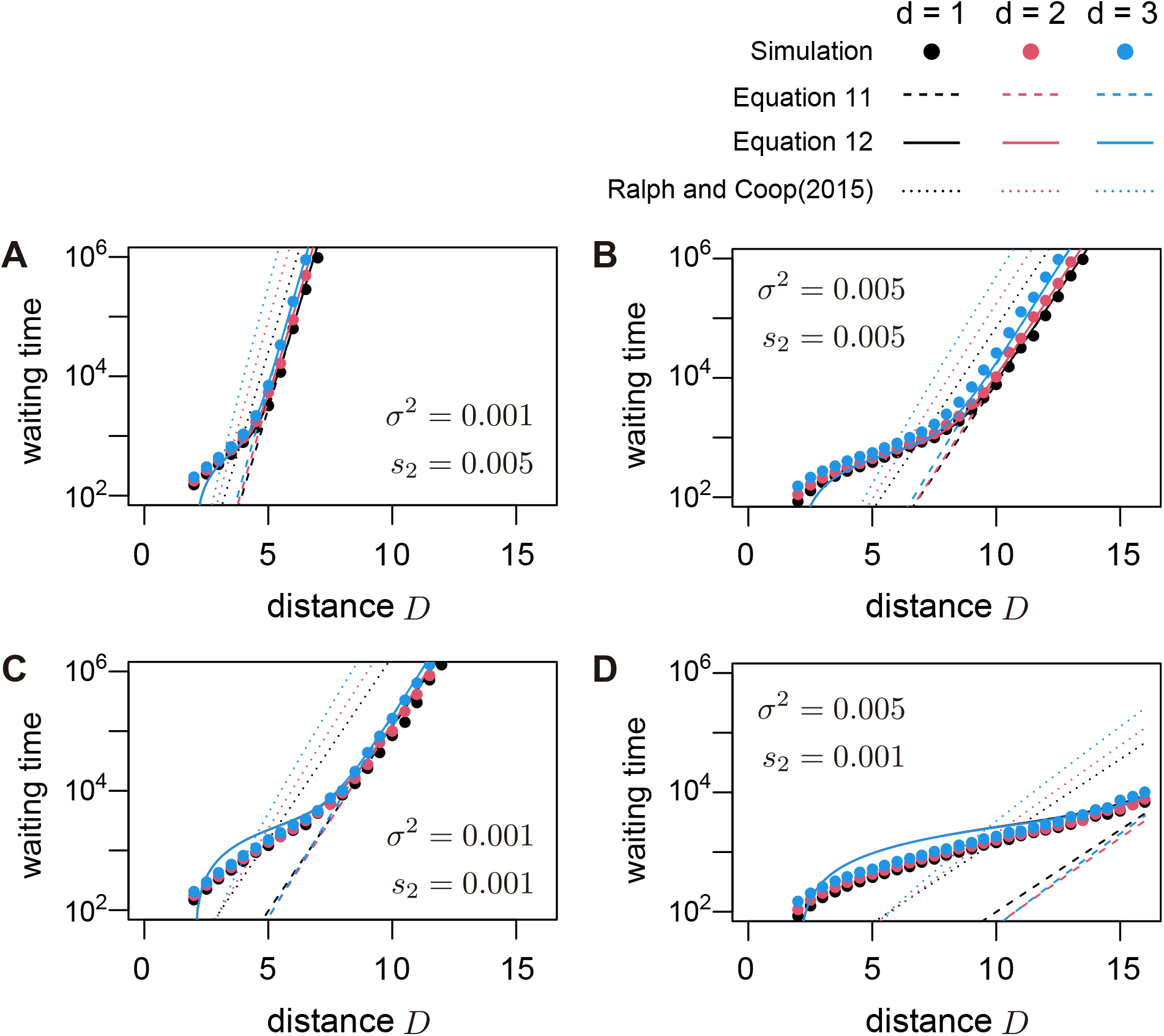
Time until local adaptation through immigration as a function of the distance between the source and target patches. Migration and selection strength in each panel corresponds to the parameters used in Figure 2. It was assumed that *s*_1_ = 0.05, *R* = 1 and *R*_*s*_ = 1. Dashed and solid lines represent the theoretical prediction calculated from Equations 11, 12, respectively. The prediction of Ralph and Coop (2015) (Equation 11) is also plotted in dotted lines for comparison. The circles show the simulation result. For each parameter set, 100 simulation replicates were run. Black, red, and blue color represent the results for *d* = 1, 2, and 3, respectively. Due to the limited range of *D*, one might think that the overall pattern is not clear in the panel D; see Figure S3 for the results for a wider range of *D*.

Although the theory works well when the population density (*ρ*) is large, the branching process approximation may break down in lower population densities because allele A may increase by random genetic drift and may no longer be rare in some areas. In such cases, the theory overestimates the waiting time (Figure B2) because local fixation of allele A reduces the extinction risk in areas where it is deleterious and promotes the establishment of each migrant lineage (see Appendix B for detailed discussion).

## Discussion

The purpose of this study is to analyze local adaptation process in the context of continuous space. Although many theoretical studies investigated the stochastic establishment process of local adaptation using two-population models (Barton 1987; Yeaman and Otto 2011; Aeschbacher and Bürger 2014; Yeaman *et al*. 2016; Tomasini and Peischl 2018; Sakamoto and Innan 2019), it is essential to extend these results to continuous space considering that actual species live in continuous space. The development of a theoretical framework based on continuous space would be increasingly important given the growing demand to fully exploit the potential of genomic data from many geographic locations (Bradburd and Ralph 2019; Battey *et al*. 2020).

This study explored the establishment of local adaptation in a multidimensional continuous space. Consistent with previous theories (Slatkin 1973; Ralph and Coop 2015), evolutionary dynamics primarily depend on the characteristic spatial scale (1/*β*_2_), which represents the ratio of selection and migrational variance. First, we investigated the establishment probability of new locally adapted mutations.

The dimensionality of space influences this process, with higher dimensions reducing the establishment probability of each allele (Figure 2). Although most alleles that ultimately establish originate within the target patch in one dimension, contributions from regions outside the target patch become increasingly important in higher dimensions (Figure 3). This pattern reflects increased mutational opportunities with greater distances in multidimensional space. Next, we examined how immigration from other patches contributes to local adaptation. Using a single lineage approximation, we derived a formula for the rate of local adaptation through this process. In this derivation, along with the number of emigrants, the probability that an emigrated allele does not return to the source patch influences the effective outflux rate. This result highlights a key difference in migration dynamics between the two-population model and the continuous space model: back migration, negligible in the former model, occurs frequently in the latter model. Notably, the present theory aligned well with the simulation results (Figure 6).

These findings enable predictions regarding the relative contributions of mutation and migration to convergent local adaptation. Based on the present results, the rate of local adaptation through mutations, *λ*_mut_, is calculated as

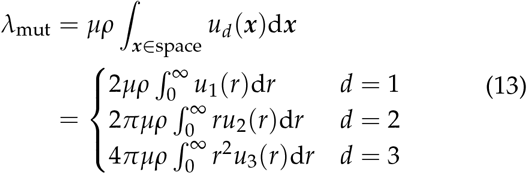

where *µ* is the mutation rate of allele A. *λ*_mig_ is given by Equation 11. Then, *λ*_mut_ and *λ*_mig_ together determine the relative contributions of each process. When both *λ*_mut_ and *λ*_mig_ are small, the waiting time until local adaptation evolves should be approximated by the exponential distribution with a mean of 1/(*λ*_mut_ + *λ*_mig_). Given the evolution of local adaptation, the probability that mutation drives this process is 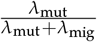. Importantly, there should exist a critical spatial distance between the source and target patches beyond which migration is highly unlikely to contribute to convergent adaptation.

As an example, assuming the same parameter values as Figure 6, we plotted how large mutation rate is required to give *λ*_mut_ = *λ*_mig_ in Figure S2. In these cases, we can predict that *D* ≈ 3–10 is a critical distance for the source patch to contribute to local adaptation through migration, assuming *µ* ≈ 10^−6^. We observed slightly larger contribution of mutation in higher dimension in Figure S2 mainly because more mutations are produced in the target patch due to its increased population size, which is given by 2*ρ* in *d* = 1, *πρ* in *d* = 2, and 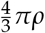 in *d* = 3 when *R* = 1. It should be noted that this pattern may not be general and depends on the parameter values assumed, especially on the sizes of the source and target patches. The strength of the present theory is its ability to predict the relative contribution over a wide parameter space.

Previous genomic analysis have revealed varying contributions of mutation and migration to convergent local adaptation across diverse species (Stern 2013; Rosenblum *et al*. 2014). An interesting case would be repeated adaptation to the freshwater in stickleback fishes. In the three-spined stickleback, many alleles are shared among freshwater populations with these alleles maintained at low frequencies in marine populations (Colosimo *et al*. 2005; Kitano *et al*. 2010; Jones *et al*. 2012), consistent with the “transporter” model (Schluter and Conte 2009), suggesting a predominant role of migration. Conversely, in the nine-spined stickleback, freshwater adaptation is caused by different variants among populations, suggesting that independent mutations play a more prominent role (Kemppainen *et al*. 2015). This difference may reflect the more limited migration in the nine-spined stickleback compared to the three-spined stick-leback (Kemppainen *et al*. 2015). Unfortunately, applying the present theory to such empirical systems remains challenging owing to the lack of spatial parameter estimates (Bradburd and Ralph 2019). However, considering the recent advancements in spatial parameter estimation methods (Ringbauer *et al*. 2017; Smith *et al*. 2023), such applications may soon become feasible in future studies. Moreover, with appropriate data, the present theoretical framework could elucidate factors that influence difference in the trajectories of convergent local adaptation among empirical systems.

## Data availability

Codes for numerical calculation and simulation are available at https://github.com/TSakamoto-evo/spatial_local_adaptation.

## Acknowledgments

I thank Sam Yeaman for his insightful comments on the earlier version. The computing resource was provided by Human Genome Center (the Univ. of Tokyo).

## Appendix A: Stepping-stone simulation

This section provides details regarding simulations conducted in *d*-dimensional space. The space is discretized into *n* segments per unit length, dividing an unit hypercube into *n*^*d*^ hypercubes, each of size 1/*n*. To achieve a migration distance variance of *σ*^2^ along each axis per generation, the migration rate between adjacent grid points is set to 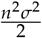. For the purpose of simulating continuous space, *n* is chosen such that the total migration per grid point, *n*^2^*σ*^2^*d*, is comparable to the selection strength (max(∼*s*_1_, *s*_2_)). In practice, *n* = 8 was assumed in *d* = 1 case, and, in cases *d* = 2 and 3, *n* = 8 was assumed for *σ*^2^ = 0.001 and *n* = 4 was assumed for 0.005. Two distinct environments are distributed across the space as defined in the Model section. Only grids where allele A is present are tracked, and the space is considered unlimited to avoid edge effects.

Each generation, allele frequencies are simulated following the Wright–Fisher model. Let *p*_*i*_ be the frequency of allele A in the *i*th grid point at generation *t*. The expected allele frequency in generation *t* + 1 is calculated as

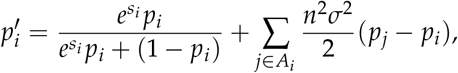

where *A*_*i*_ represents the set of grids contiguous to the *i*th grid (von Neumann neighbourhood). In grids where both environments coexist, the selection strength *s*_*i*_ is determined by the weighted average of *s*_1_ and −*s*_2_. Specifically, if the *i*th grid consists of *x* proportion of environment 1 and 1 − *x* proportion of environment 2, then *s*_*i*_ is determined as

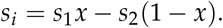

The number of allele A in the next generation is then deter-mined following Binom 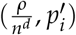.

Establishment of allele A is recorded when its frequency exceeds 0.5 within the target patch. Specifically, the establishment is recorded when the following equation is satisfied:

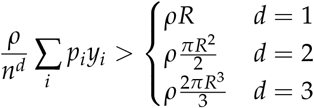

where *p*_*i*_ is the frequency of allele A in the *i*th grid and *y*_*i*_ is the proportion of the area belonging to the target patch within the *i*th grid.

## Appendix B: Simulation results for lower population density

In the main text, the simulation results are presented where large *ρ* (*ρ* = 2^16^) is assumed for comparison with the theoretical prediction (Figures 2, 6). The large *ρ* assumption was made because the branching process approximation used in the derivation is valid when the population density is sufficient large to prevent local fixation of deleterious alleles. In this section, we run simulations assuming lower population densities to examine what patterns are observed and how the theory deviates from simulation results at lower *ρ*. We first focused on the establishment probability of a new mutation in Figure B1. Using the same parameter set as in Figure 2, we assume four different *ρ* (*ρ* = 2^10^, 2^12^, 2^14^ and 2^16^) for *d* = 1, 2, and three *ρ* (*ρ* = 2^12^, 2^14^ and 2^16^) for *d* = 3. The results for *ρ* = 2^10^ are not provided for *d* = 3 because the population size of each grid becomes too small and the simulation is infeasible for some cases (see also APPENDIX A). Figure B1 shows that the establishment probability observed in the simulation increases slightly in some cases. Such an increase was observed for *d* = 1, 2 with *ρ* = 2^10^ and *σ*^2^ = 0.001, and for *d* = 3 with *ρ* = 2^12^ and *σ*^2^ = 0.001. This increase of the establishment probability is due to the local fixation of allele A in some unfavored areas (i.e., environment 2), which reduces the effectiveness of selection.

We next examined the waiting time until immigration fuels local adaptation in Figure B2. Figure B2 shows that our theory generally works well across the different population density. We observe that the theory overestimates the waiting time for parameters where the establishment probability is higher than the theoretical prediction, consistent with Figure B1.

We finally examined whether the density at which the branching process approximation breaks down is relevant to the analytical prediction. Wright’s neighborhood size *N*_loc_ is calculated for this purpose because it quantifies the effect of random genetic drift in continuous spaces (Wright 1946). For each dimension, *N*_loc_ is given by

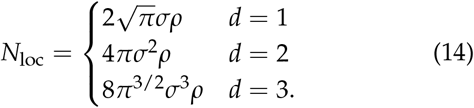

The value of *N*_loc_ for each parameter set is provided in Figure B1. Unfortunately, the relationship between *N*_loc_ (or *N*_loc_*s*_2_) and the breakdown of the approximation is unclear. This pattern suggests that the validity of the branching process approximation depends on the population density, selection strength, migrational variance, and spatial dimension in a complex manner. Delineating this relationship would be the subject of future theoretical studies.

**Figure B1.**
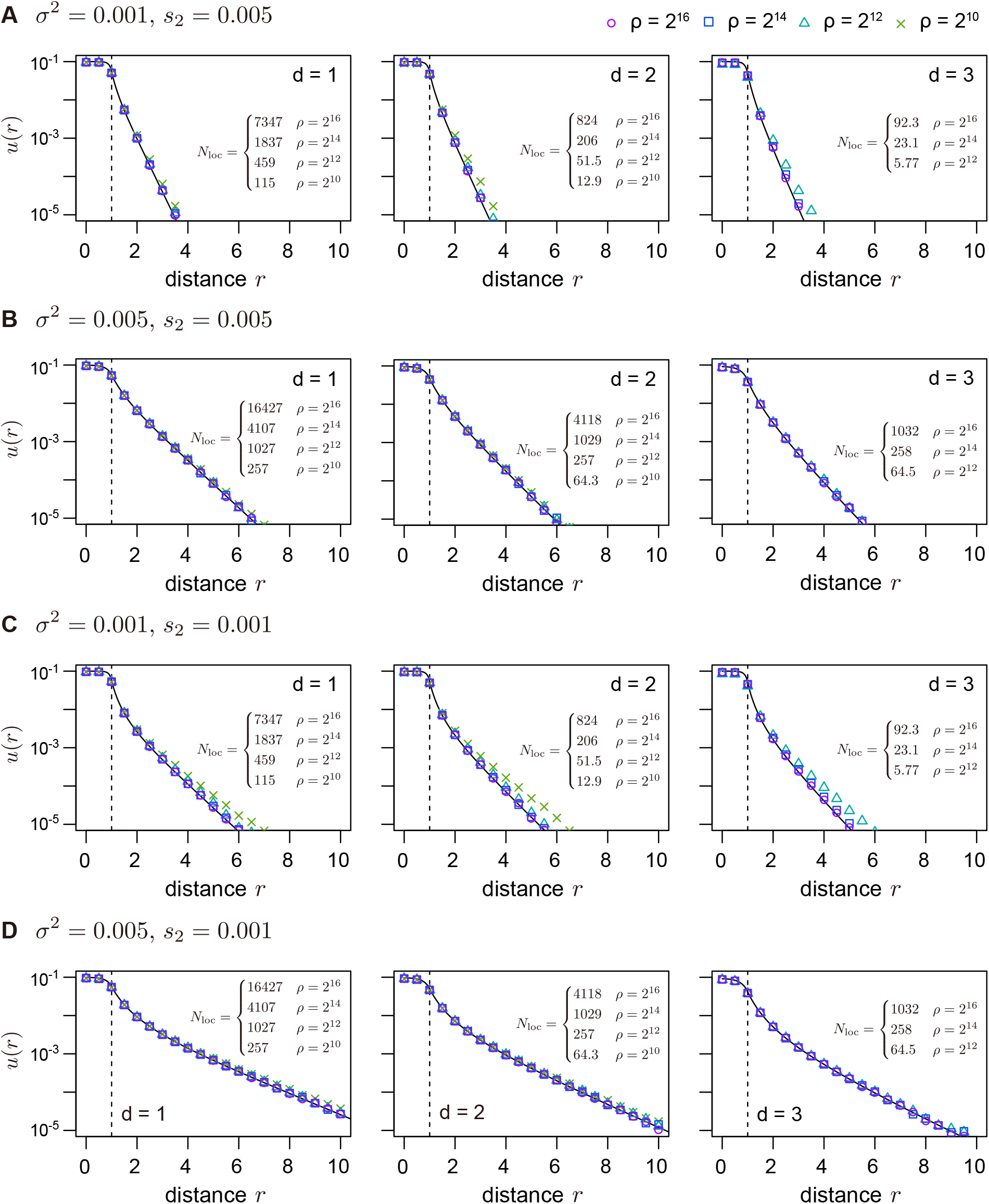
Establishment probability a locally adaptive mutation in lower population densities. Parameter values mirror those given in Figure 2 except for population density. For each parameter set, 10^7^ simulation replicates were run. Vertical dashed lines are drawn at *r* = *R*, marking the boundary between the two environments.

**Figure B2.**
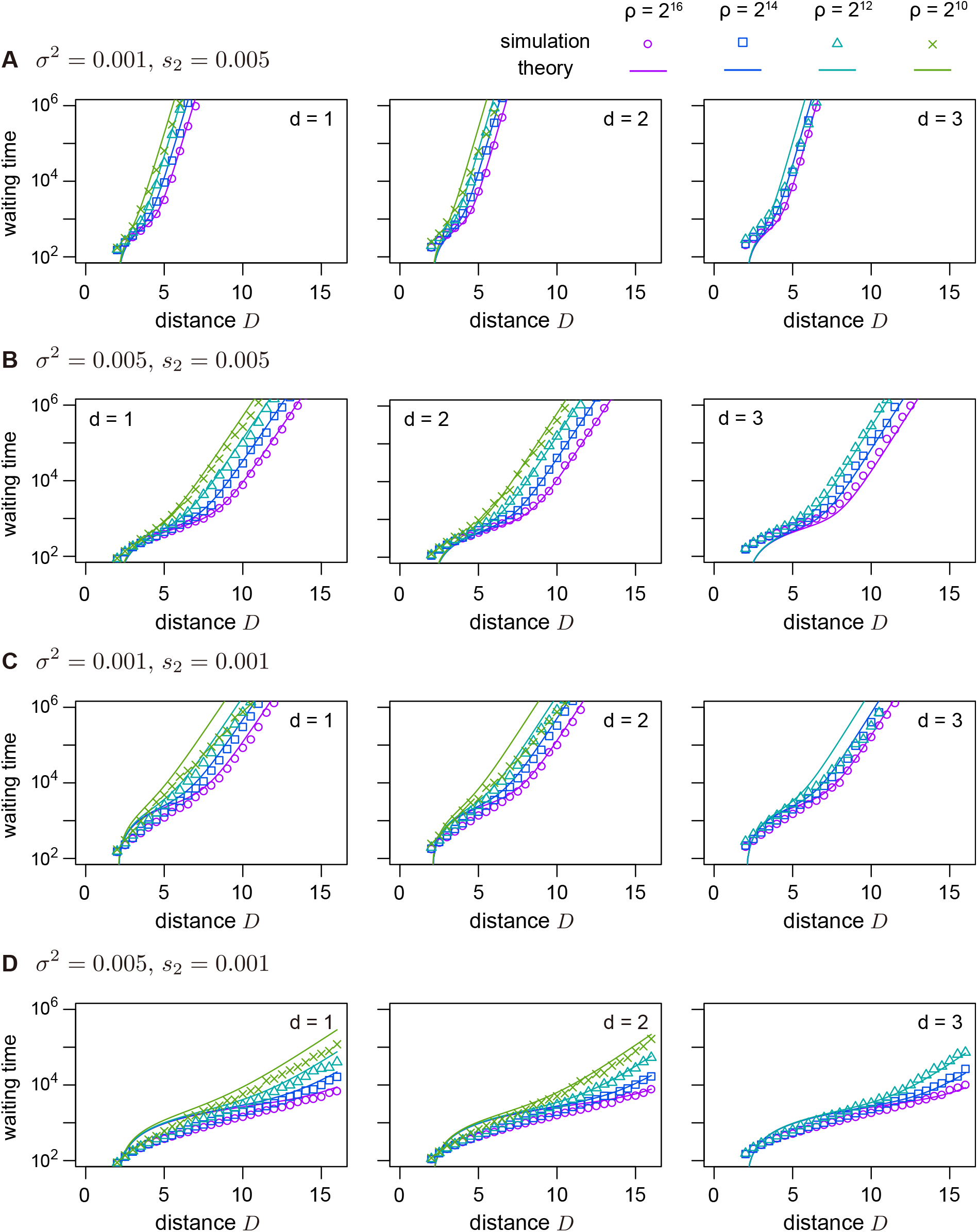
Waiting time until local adaptation through immigration in lower population densities. Parameter values mirror those given in Figure 6 except for population density. Each color represents the results for different population density. For each parameter set, 100 simulation replicates were run.

**Figure S1.**
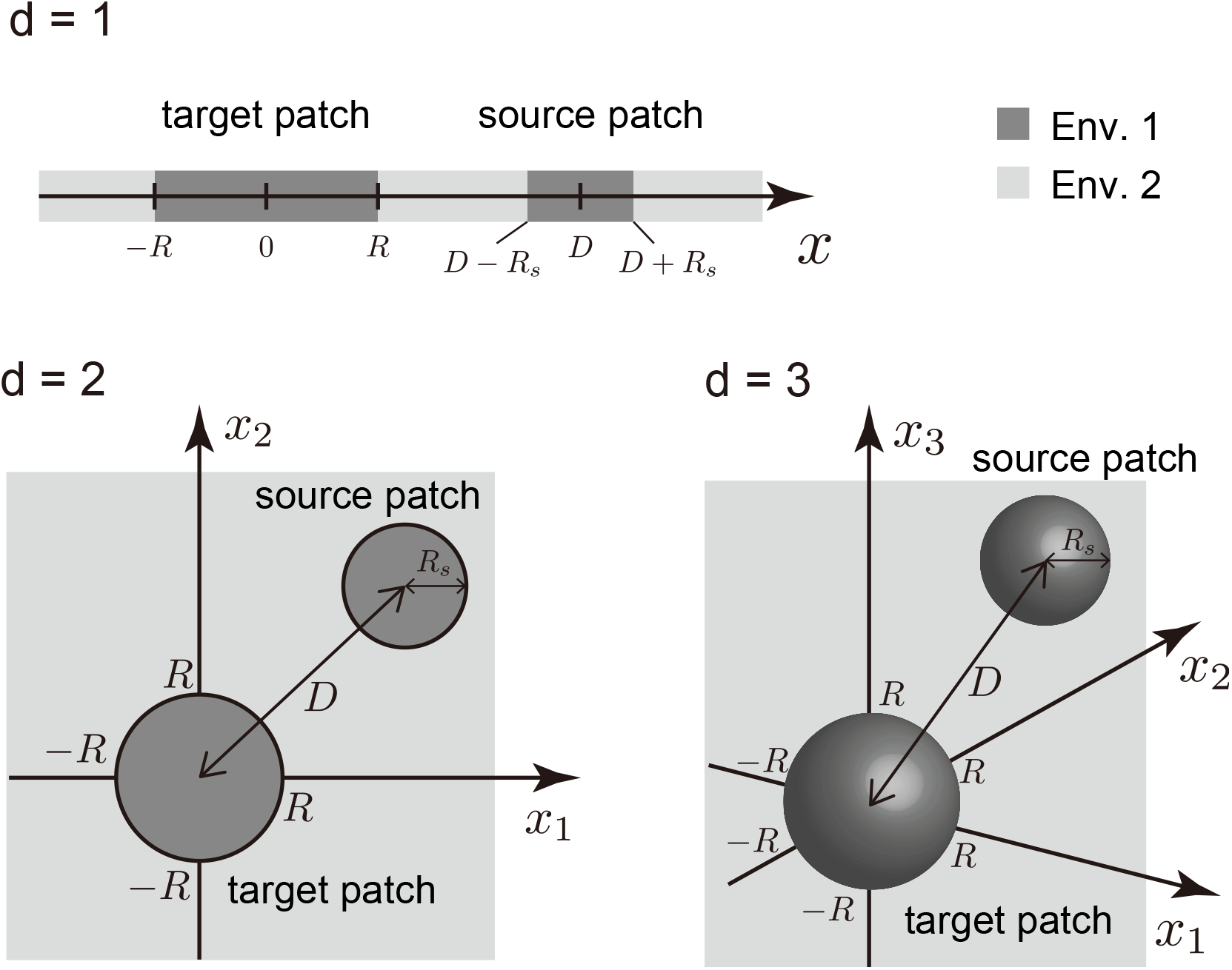
Illustration of the two-patch model.

**Figure S2.**
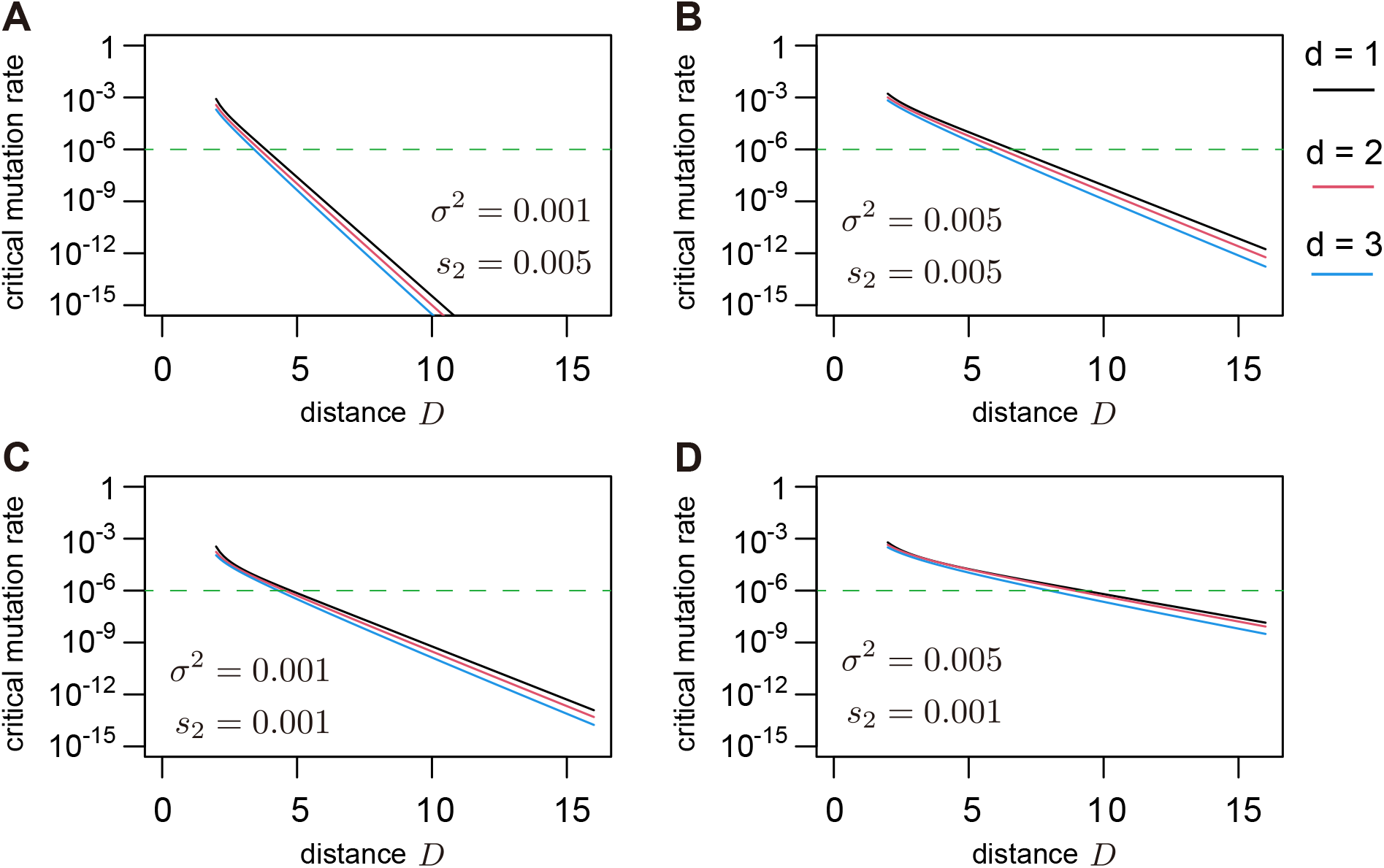
Mutation rate at which the relative contributions of mutation and migration are balanced (*λ*_mut_ = *λ*_mig_). Green dashed lines are drawn at *µ* = 10^−6^. Parameter values mirror those given in Figure 6.

**Figure S3.**
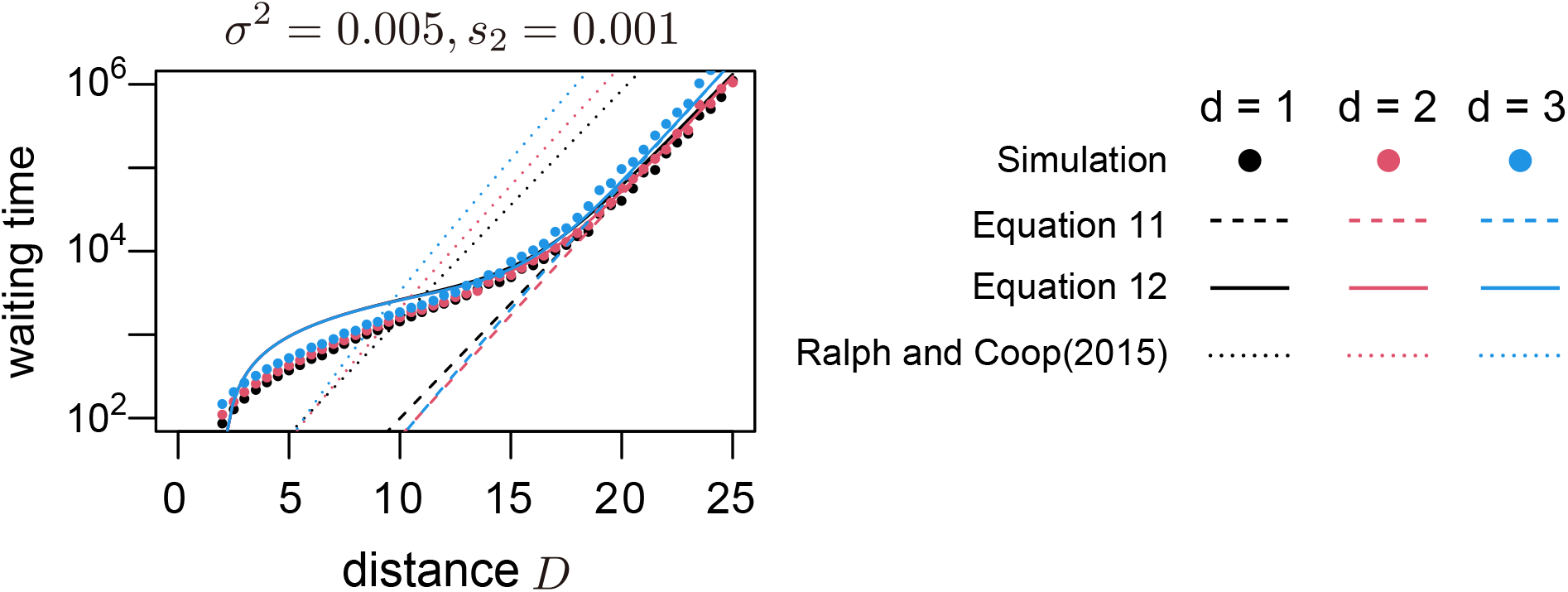
A version of the panel D of Figure 6 assuming a wider range of *D*.

